# Individual-to-individual mapping of color similarity structures: Is my “red” relationally your “red”?

**DOI:** 10.64898/2026.02.13.705699

**Authors:** Yu Togashi, Yuko Yotsumoto, Chihiro Hiramatsu, Naotsugu Tsuchiya, Masafumi Oizumi

**Affiliations:** Graduate School of Arts and Science, The University of Tokyo, Meguro-ku, Tokyo 153-0041, Japan; Faculty of Design, Kyushu University, 4-9-1 Shiobaru, Minami-ku, 8158540, Fukuoka, Japan; School of Psychological Sciences, Monash University, Clayton, VIC 3168, Australia; Computational Neuroscience Laboratories, Advanced Telecommunications Research, Seika-cho, Kyoto 619-0288, Japan; Theoretical Sciences Visiting Program (TSVP), Okinawa Institute of Science and Technology Graduate University, Onna-son, Oki

**Keywords:** qualia, qualia structure, consciousness, color vision, unsupervised alignment

## Abstract

Is my “red” the same as your “red”? Although we cannot directly test whether two individuals’ experiences (qualia) are identical, we reframe this question as an empirical one: whether my “red” is relationally your “red”. We addressed this question by finding the best possible mappings between color similarity structures across individuals, without using color labels. We collected a complete set of similarity judgments across 93 colors from each of 11 participants and compared their structures. These comparisons revealed two groups of four participants in this sample whose structures aligned robustly within each group: the best possible mappings matched the same color stimuli, such as my “red” to your “red”. Between groups, however, the best mappings did not match the same stimuli. The remaining participants showed structural variations that did not fit either group. This framework opens a path to classifying diverse color experiences through their relational structures.

## INTRODUCTION

From childhood on, many of us wonder about a fundamental question: Are our subjective experiences, such as of color, the same as those of others? For instance, is my “red” the same as your “red”? Traditionally, the qualitative aspects of consciousness, or qualia for short^1,2^, have been considered unaddressable within the scientific framework due to their inherent privateness and ineffability^3,4^. Even if two people agree that an apple is red, there is no guarantee that their internal experiences are identical^5– 7^. The same apple that appears red to you may appear green to another person, as the longstanding philosophical argument of “inverted spectrum”^5^ posits.

Because it is difficult to characterize the quality of an experience in isolation, one promising approach to deal with qualia quantitatively is to focus on relational structures among qualia^8–25^ (Figures 1A and 1B). Rather than referring to absolute qualities, it is feasible to report the relationships between them, such as similarities: e.g., “red is similar to orange” or “red is dissimilar to green” (Figure 1A). By aggregating a comprehensive set of such relational reports, we can objectively characterize and quantify the relational properties of the underlying subjective structures (Figure 1B). We call such relational structures “qualia structures”^24^. We note that relational reports such as similarity judgments may in part reflect factors other than color experience itself, such as color categories, lexical knowledge, or associations with familiar objects. Therefore, an empirically estimated similarity structure should be considered as an approximation of the underlying qualia structure. In what follows, we use “color similarity structure” to refer to the empirically estimated structure, and reserve “qualia structure” for the hypothesized experiential structure that it approximates.

**Figure 1.**
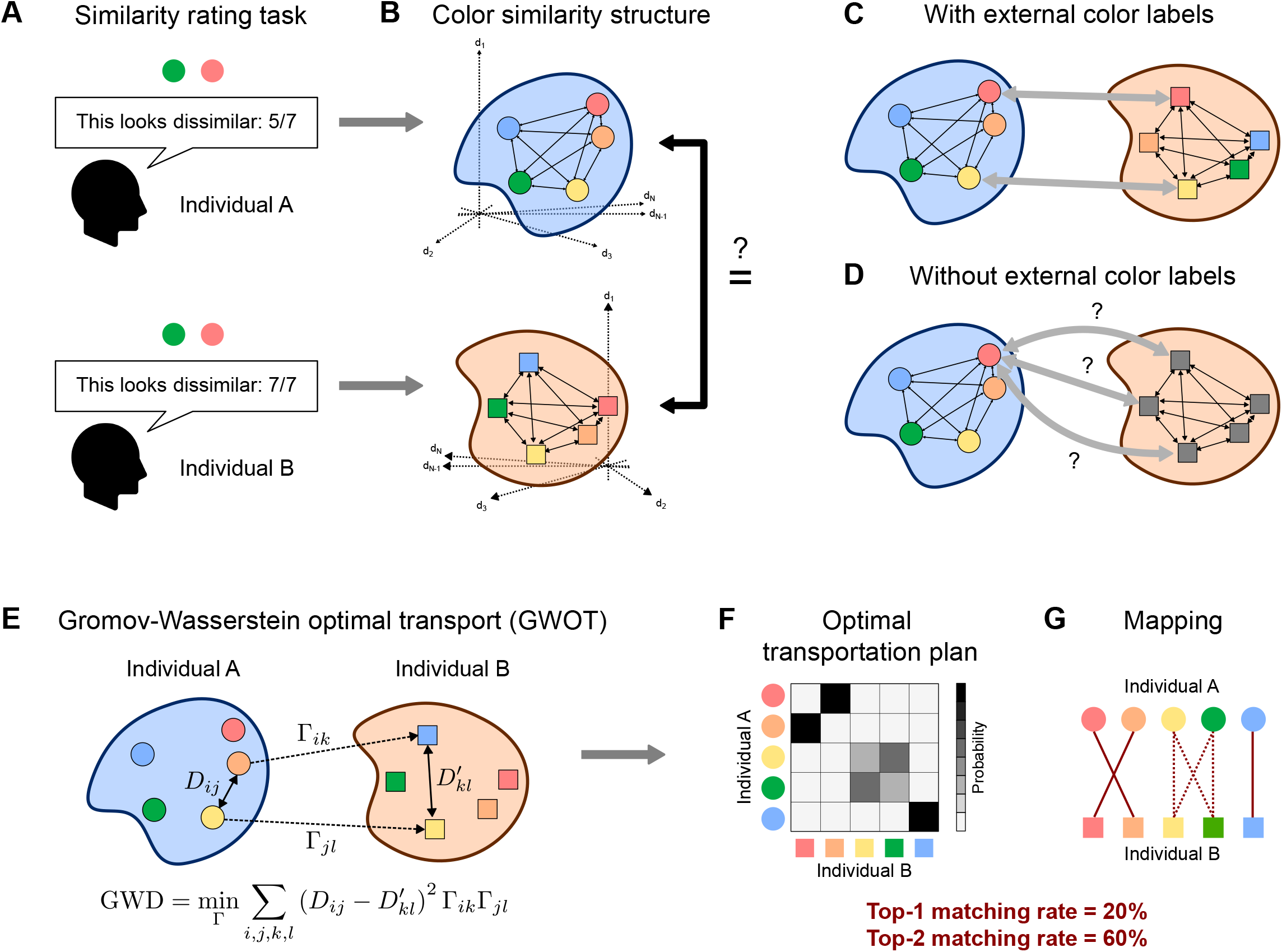
Overview on one-versus-one unsupervised alignment between individual color similarity structures. (A) To estimate the color similarity structure for each individual, we collected a complete dataset of 4,371 pairwise similarity judgments for 93 colors from individuals. (B) Color similarity structures are estimated for each individual by applying multidimensional scaling (MDS) to the reported similarity ratings. (C) Supervised alignment of color similarity structures, which assumes a single mapping externally determined by color labels. (D) Unsupervised alignment of color similarity structures, which finds the optimal mapping itself based solely on their internal geometries. (E) Gromov-Wasserstein optimal transport (GWOT). The elements of *D* and *D′* are the distances (dissimilarities) between the color experiences for each individual. Γ is the transportation plan, which designates the correspondence between color similarity structure. GWOT finds the optimal correspondence between two structures that preserves the distance relationship between structures as much as possible. (F) Example of the optimal transportation plan Γ, where the (*i, j*) element represents the probability that color *i* in one individual is mapped to color *j* in the other. From the optimal transportation plan, the degree of correspondence between the two structures is quantified as the top-*k* matching rate (see STAR Methods). (G) The optimal transportation plan can be seen as a mapping between two structures, which allows for coarse, many-to-many mapping (represented as dashed lines).

Although this structural approach has gained substantial momentum in consciousness science^21–23,25^, its core empirical question has remained unresolved: do two *specific* individuals actually share the same relational structure of experience? This question is subtle because of a classic philosophical challenge, the inverted spectrum^5–7^: even if two people’s color qualia structures appear identical, one person’s color experiences could be covertly swapped relative to the other’s, i.e., my red might be your green, while every reported similarity relation remains intact. Palmer^9^ scientifically sharpened this concern for color, arguing that in an idealized, symmetric model of color space a red–green swap preserves all similarity relations and therefore escapes behavioral detection, so that no accumulation of similarity judgments could establish whether your red is my red. Whether such a structure-preserving swap truly escapes detection once color experiences are densely and empirically sampled, however, has never been tested. Addressing this requires what has so far been missing: (1) a method to compare the color similarity structures of two individuals in a logically principled way that directly confronts this inverted-spectrum possibility, and (2) the empirical construction of individual-level color similarity structures rather than group averages.

Meeting the first requirement calls for an unsupervised approach that compares two color similarity structures without presupposing any correspondence between individuals’ experiences^24,26^. This is essential because a method that begins by pairing my “red” with your “red” would simply assume away the inverted-spectrum possibility rather than test it. However, conventional methods to compare similarity structures have primarily relied on such supervised methods^27–31^, which assume an externally fixed correspondence between the two structures, as is typical in Representational Similarity Analysis (RSA)^32,33^ (Figure 1C): they presuppose that my “red” qualia maps to your “red” qualia and then quantify structural agreement under this presumed mapping. Instead, our unsupervised alignment method discovers the optimal mapping between two color similarity structures from their internal geometries alone (Figures 1D–1G). This optimal mapping is the maximally structure-preserving mapping^34^: the mapping that preserves the distance relationships among qualia across individuals to the greatest degree. In concrete terms, such a mapping sends two colors that are similar for one individual to two colors that are similar for the other individual, and likewise for dissimilar colors. By recovering this mapping without color labels, we can determine whether your “red” relationally corresponds to my “red” (a stimulus-matched correspondence) or to my “green” (an inversion).

It is worth noting that our aim is to reframe the original question of whether your “red” is the same as my “red” into a specific, empirically answerable one, namely whether your “red” is relationally similar to my “red”. What we do within this framework is to search for the best mapping between two structures among all possible mappings, without using color labels. When the mapping that emerges as the optimal one is stimulus-matched (e.g., red-to-red), other mappings that involve a swap (e.g., a red-to-green swap) become less plausible accounts of how the two individuals’ color experiences relate. Whether the two experiences matched across individuals are phenomenally identical, however, is a further question that our framework alone cannot address.

While we already applied this framework to inter-group comparison of color similarity structures^24^, that study left a critical question unanswered: do color similarity structures actually align across specific individuals? Using color as a model domain, we previously demonstrated that averaged color similarity structures align within color-neurotypical and atypical groups. However, a group-level analysis cannot determine whether structures are shared or divergent between any two specific people. Statistical averaging tends to converge toward a common mean, which may obscure crucial individual differences in qualia structures. This concern also applies to color vision, where well-documented diversities in both neurophysiological and perceptual properties exist even among color-neurotypical populations (for review, see Bosten^35^). Consequently, testing whether structures of color experiences are shared or divergent between two specific individuals requires a direct, individual-to-individual comparison.

Here, we performed the first direct, one-versus-one alignment of individual color similarity structures. Using color experience as a test case and encompassing participants with diverse forms of color vision, we asked whether structure-preserving swaps such as a red–green inversion truly escape detection, and more broadly, whether our framework can recover the structural correspondence and divergence of subjective experience between specific individuals. To achieve this, we collected a complete set of 4,371 pairwise similarity judgments (Figure 2A) for 93 colors (Figure 2B) from 11 participants, for about 10 hours spread over 5 days per participant (Figure 2C). Participants were assessed for their (a)typicality of color vision based on both self-report and the Farnsworth–Munsell 100 Hue Test, which assesses local discrimination sensitivity (see STAR Methods)^36^ (Figure 2D). This rich dataset enables the first direct, individual-to-individual comparison of color similarity structures across a broad and densely sampled color space, allowing for a rigorous evaluation of how color experience varies among people. Our primary finding is that we identified two groups of participants whose color similarity structures can be robustly aligned in an unsupervised manner. Within each group, structures aligned in a stimulus-matched manner—my “red” is relationally your “red”—disfavoring the plausibility of structure-preserving swaps such as red-to-green. Between the two groups, by contrast, structures did not align, indicating cases where my “red” is not relationally your “red”. Coexisting with these clusters, however, we observed nuanced variations in color similarity structures among certain participants, which revealed perceptual diversity not fully captured by the conventional hue test. These findings provide an empirical foundation for a structure-based taxonomy of color experience grounded directly in its phenomenology, while also capturing fine-grained experiential diversity that defies simple categorization. Although we focus on color, where perceptual diversity is well-documented and the philosophical stakes are sharpest, our one-versus-one alignment framework is inherently domain-general and applicable wherever similarity judgments can be collected across a broad stimulus space.

**Figure 2.**
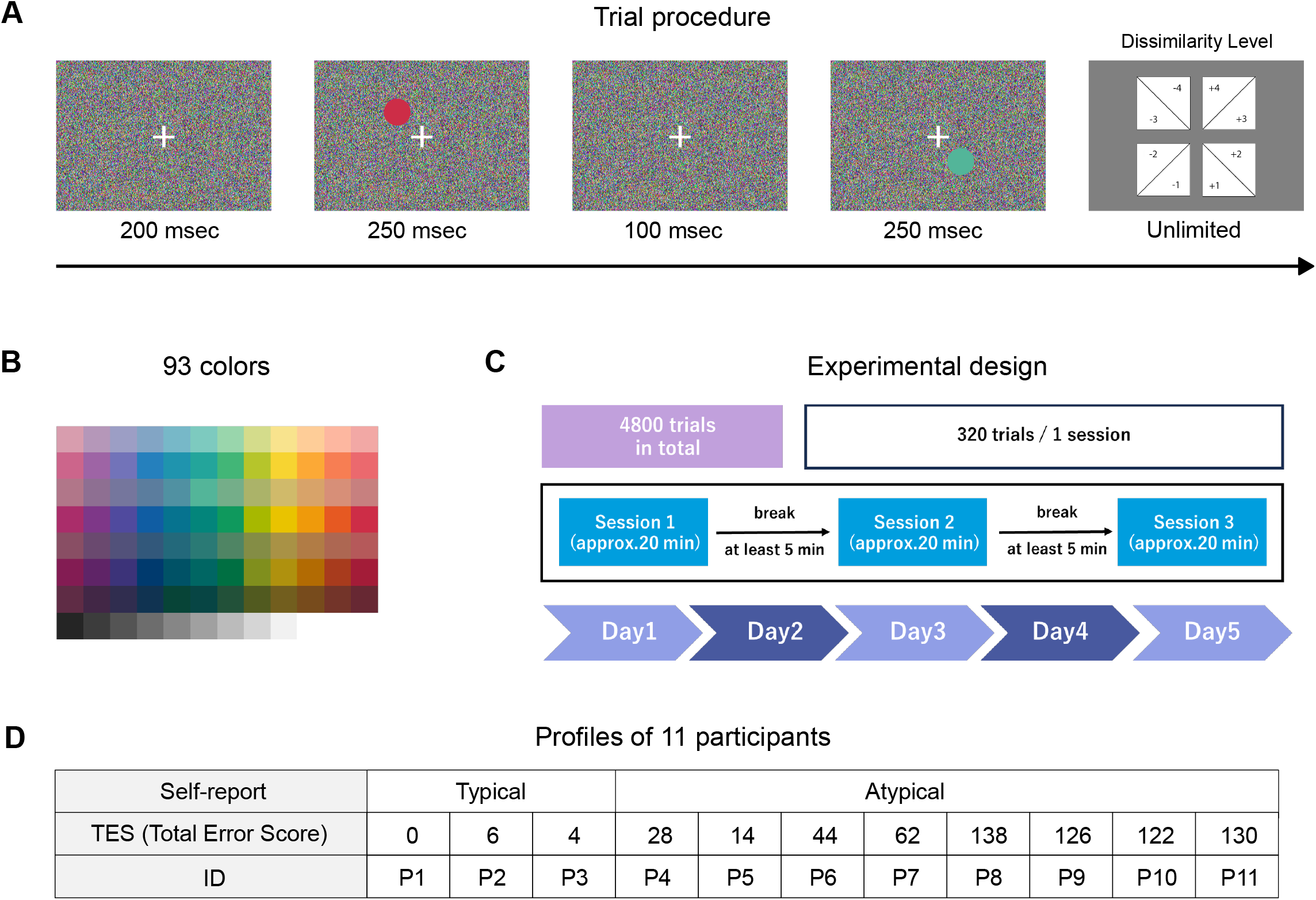
Experimental procedure and participants demography. (A) Schematic of the main trial procedure (for details, see STAR Methods). (B) 93 colors used in our study. (C) Experimental design of the similarity rating task. Each participant went through a complete set of similarity rating tasks for about 10 hours spread over 5 days (for details, see STAR Methods). (D) Profiles of 11 participants. Participants were assessed for their (a)typicality of color vision based on self-report and the FM100 hue test. A higher TES (Total Error Score) indicates lower color discrimination performance on the FM100. For detailed error patterns, see Figure S1.

## RESULTS

### Overview of one-versus-one unsupervised alignment between individual color similarity structures

Our investigation is guided by two primary objectives. First, we determine whether color similarity structures can be empirically aligned between at least one pair of individuals in a purely unsupervised manner. This is a non-trivial starting point, as there is no prior guarantee that a shared qualia structure exists between any two people given known perceptual diversities. Second, we investigate whether there exists a group of individuals where structures align consistently among members. Such a finding would allow us to classify individuals based on the global structure of their experiences rather than on local sensitivities assessed through performance on standard discrimination tests. If successful, these alignments would suggest that the relational geometry among qualia is shared within specific populations, providing an empirical approach to the long-standing philosophical puzzle, “Is my red your red?”^24^

To empirically identify the inter-subjective correspondence of color similarity structures, we searched for the optimal mapping between two individuals’ color similarity structures using Gromov-Wasserstein optimal transport (GWOT)^34^. This method identifies the mapping that best corresponds to the experiences of two individuals by minimizing the Gromov-Wasserstein Distance (GWD), which quantifies the distortion of distance relationships between two structures. The result is the optimal transportation plan Γ (Figure 1F), in which each element Γ_*ik*_ represents the probability that the *i*-th color experience in one person corresponds to the *k*-th color experience in the other. The optimal transportation plan can be seen as a relaxed mapping that allows for coarse many-to-many correspondence (as shown in Figure 1G), and effectively create a translation dictionary between the qualia of two different people. This mapping is deemed “optimal” because it represents the most compatible alignment of two similarity structures among all possible graded mappings. Once this optimal mapping is established, we can examine whether your “red” relationally corresponds to my “red” (Figures 1F and 1G). We quantify the degree of this correspondence using the top-*k* matching rate, which measures the probability that colors are consistently mapped across individuals (see STAR Methods).

### Complete collection of similarity judgments for 93 colors from 11 participants

To perform one-versus-one unsupervised alignment across individuals, we collected a complete dataset of 4,371 pairwise similarity judgments for 93 colors from 11 participants (Figure 2A; see STAR Methods). The colors were selected from the Practical Color Coordinate System (PCCS)^37^, following the stimulus set used by Saji et al.^38^. The final set comprised 12 hues sampled across 7 tones, along with 9 achromatic colors ranging from white to black (Figure 2B). Given that completing 4,371 judgments requires substantial time and mental effort, we distributed the experiment over five days, with approximately two hours of data collection per day (Figure 2C).

11 participants (P1–P11) were assessed for the (a)typicality of their color vision based on both self-report and the Farnsworth–Munsell 100 Hue Test (FM100) (Figure 2D). In our recruitment, we explicitly stated that we sought participants with both typical and atypical color vision, with a particular emphasis on recruiting individuals with atypical color vision. During the screening process, prospective participants were asked to self-report whether they were aware of having typical or atypical color vision using a two-alternative forced-choice question. Based on these self-reports, we recruited 3 participants with self-reported typical color vision (P1–P3) and 8 participants with self-reported atypical color vision (P4–P11). In addition to self-reports, all participants completed a standard color vision assessment, the Farnsworth– Munsell 100 Hue Test (FM100)^36^. Performance on the FM100 is summarized by the Total Error Score (TES), with higher scores indicating lower color discrimination (Figure 2D). As a reference point, it has been reported that, among color-neurotypical individuals, approximately 16% score TES 16, while another 16% score TES > 100; these norms are based on a population aged 15 to 45, which encompasses the age range of our participants (see STAR Methods)^39^. Note that the TES on FM 100 alone does not allow for a definitive distinction between typical and atypical color vision. Detailed error patterns on FM100 for all participants are shown in Figure S1.

### Unsupervised alignment reveals robustly aligned groups of shared color similarity structure

By applying the unsupervised alignment method to all pairwise combinations of participants, we identified two candidate groups of individuals in the present sample, each of which showed robust within-group alignment. While we visualize representative alignments below, a comprehensive analysis of all pairwise combinations is provided in Figure S2. The first group, which we refer to as Group A, consisted of P1, P2, P3, and P4. The second group, Group B, comprised P8, P9, P10, and P11, and showed equally robust alignment. These groupings emerged solely from the alignment of similarity structures, without reference to self-report or FM100 performance; their correspondence to these assessments is examined in a later section.

We first visually demonstrate the results of unsupervised alignment within Group A. Figure 3A shows the estimated color similarity structures of P1–P4, where the distance between two colored points reflects the subjective dissimilarity of these two color experiences for each participant. After obtaining the color similarity structures of each participant, we performed GWOT to find the optimal transportation plan Γ between each pair. Using this plan, we rotated and flipped the color similarity structures of P1, P2 and P4 and overlaid them onto that of P3 to align them as much as possible (see STAR Methods and Video S1) (Figure 3B). Crucially, this entire alignment procedure was performed in a purely unsupervised manner without relying on color labels. We emphasize this fact by rendering all points in Figure 3B as black. Subsequently, to evaluate the correspondence (Figure 3C), we colored each point according to its physical stimulus label (Figure 2B). This visualization intuitively captures the robust alignment (i.e., top-1 matching rate of 48%) among these four participants given that points of similar color, which correspond to each participant’s qualia evoked by the color stimuli, cluster tightly together in this common representation space.

**Figure 3.**
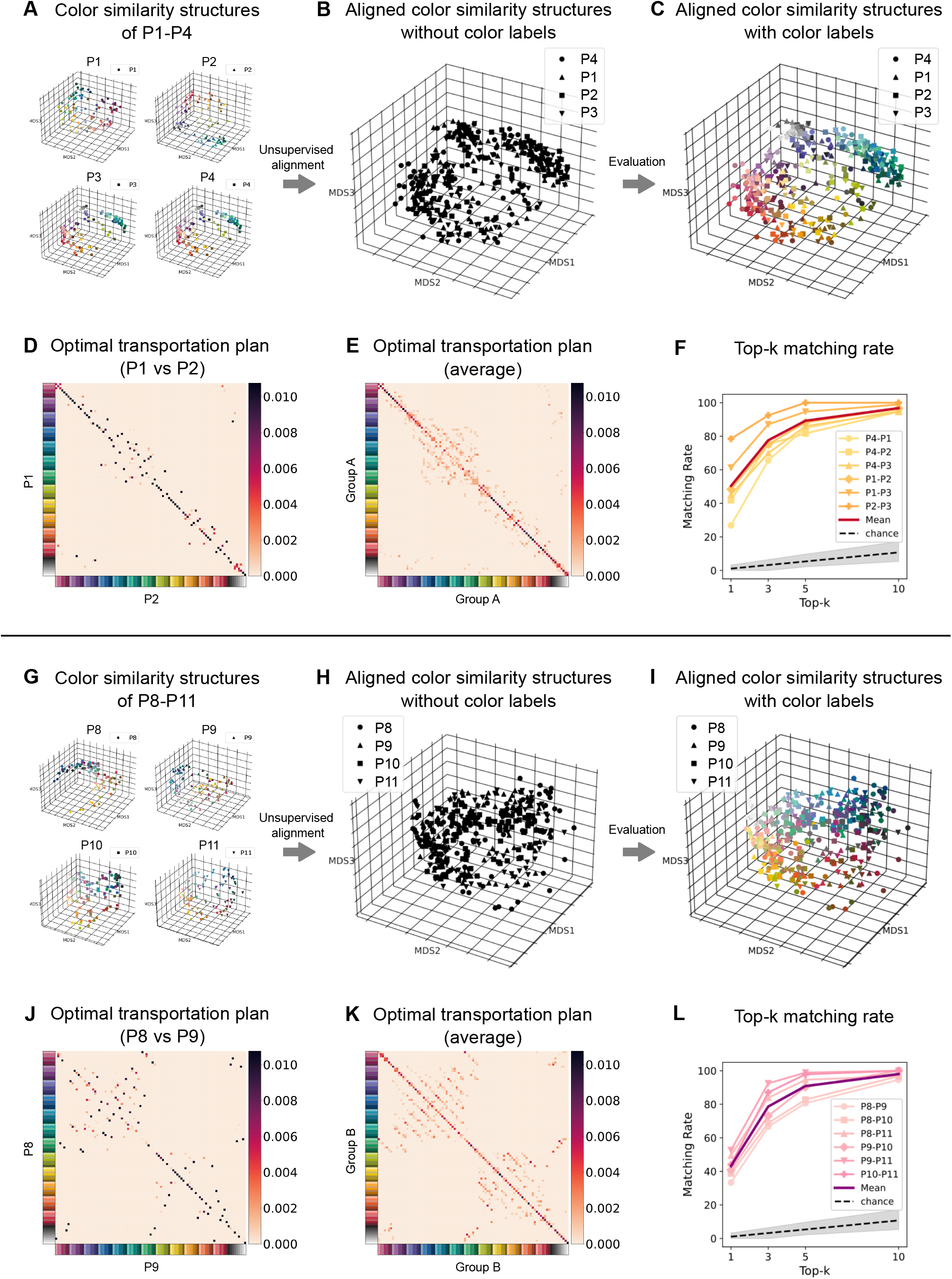
Individuals form two robustly aligned groups. (A) Color similarity structures of four participants, P1, P2, P3, and P4. Structures were first estimated by applying 5D multidimensional scaling (MDS) to individual similarity ratings. For visualization, we reapplied 3D MDS to the resulting 93 × 93 distance matrices (see STAR Methods). Axes represent the MDS dimensions. Each point represents a color experience, and the Euclidean distance between points reflects the perceived dissimilarity between the two colors. (B) Aligned color similarity structures of P1–P4. Color similarity structures of P1, P2 and P4 are overlaid onto P3’s color similarity space, using the optimal transportation plan Γ (see STAR Methods). Note that the alignment procedure is performed in a purely unsupervised manner without reliance on color labels. (C) Aligned color similarity structures of P1–P4, colored with the external color labels. (D) Optimal transportation plans Γ between P1 and P2. Rows and columns correspond to the 93 colors of two individuals, respectively. The (*i, j*) element shows the probability that color *i* in one individual is mapped to color *j* in the other. (E) The average optimal transportation plan over all pairs of P1–P4 (n = 6 pairs), calculated as the average of all the optimal transportation plans and their transported matrices. (F) Top-*k* matching rate for all pairs of P1–P4 (n = 6 pairs). For each pair, the top-*k* matching rate is plotted against *k* (1, 3, 5, and 10). The highlighted line represents the mean matching rate across pairs. The dashed line represents the mean chance-level matching rate, and the gray-shaded region represents its 95% percentile interval, estimated from 10,000 random transportation plans (see STAR Methods). (G–L) Same format as (A)–(F), but for the four participants P8, P9, P10, and P11 (n = 6 pairs in K and L).

Examination of the optimal transportation plans Γ (Figure 3D) reveals the fine-grained nature of this structural correspondence, and confirms a high degree of consistency in color mappings across individuals (P1–P4). The example optimal transportation plan (Figure 3D: P1 vs P2) exhibits high values along many diagonal elements, indicating that many colors are mapped to the same colors across individuals, i.e., “red” for one individual is relationally “red” for the other, not “green” or “blue”. This trend is confirmed by the average optimal transportation plans Γ across all pairs of P1–P4 (Figure 3E), where most diagonal elements clearly show high values. This tendency is reflected in consistently high matching rates across all pairwise combinations of P1–P4 (Figure 3F), all of which fall outside the 95% percentile intervals of the chance-level estimated by generating random transportation plans (see STAR Methods). For example, in the alignment between P1 and P2, the top-1, -3, -5, and -10 matching rates are 48%, 75%, 88%, and 97%, respectively, compared with chance-level expectations of 1.1%, 3.2%, 5.4%, and 10.7%. On the other hand, the optimal transportation plans (Figures 3D and 3E) exhibit off-diagonal values around bluish and greenish hues, revealing systematic confusions in these color ranges. These swapped mappings likely arise from the subjective resemblance among these colors (Figure 3A). Taken together, these color similarity structures robustly align within this specific subset of participants, namely Group A, while allowing slight divergence around bluish and greenish hues.

Similarly, we observed robust unsupervised alignment within Group B (P8–P11). We found that the structures of P8–P11 (Figure 3G) could be overlaid into a common space with high precision (Figures 3H and 3I). Overall, the same or similar colors from different individuals were located close to each other, although some confusion between dissimilar colors was also observed (Figure 3H). The optimal transportation plan Γ (Figures 3J and 3K) further reveals detailed aspects of this structural correspondence. The matching rate calculated from the optimal transportation plans falls outside the 95% percentile interval of the chance-level across all pairs (Figure 3L), indicating the shared geometry in their color space. For example, in the alignment between P8 and P9 (Figure 3J), top-1, -3, -5, and -10 matching rates are 33%, 67%, 80%, and 95%, respectively. Notably, however, swapped mappings were observed in certain color ranges, for example among purplish, greenish, and achromatic hues (Figure 3J). These mismatch patterns were consistently observed across all pairs of participants (Figure 3K). These colors are subjectively similar for these participants (Figure 3G), and this similarity is reflected in the optimal transportation plan as such swapped mappings (see the case of the Group A in Figure 3E for comparison). Taken together, these results indicate that color similarity structures can robustly align this specific subset of participants, namely Group B.

Finally, we confirmed that these emerging groups are structurally distinct by demonstrating that they do not align robustly with one another. In contrast to the robust within-group consistency, the color similarity structures fail to align between Group A (P1–P4) and Group B (P8–P11) (Figure 4A) in a stimulus-matched manner, indicating that “red” for one individual is not “red” for the other. For example, the optimal transportation plan between P2 and P9 (Figure 4B) contains many off-diagonal elements, and its matching rate (top-1: 1.1%, top-3: 1.1%, top-5: 3.2%, top-10: 6.5%) is essentially at chance (top-1: 1.1%, top-3: 3.2%, top-5: 5.4%, top-10: 10.7%), reflecting substantial cross-mappings across a wide range of colors. Notably, however, some individual pairs (e.g., P1 vs. P8) exhibit moderate alignment (Figures 4C and 4D). Although their matching rates (top-1: 7%, top-3: 11%, top-5: 17%, top-10: 36%) remain below those observed within either group, they nonetheless exceed the 95% percentile interval of the chance-level, indicating coarse alignment across the two groups. These coexisting features of strong divergence and moderate alignment are systematically captured in the matching rates across all Group A-B pairs (Figure 4E). These results indicate that while the two groups possess distinct color similarity structures, they retain broad geometric commonalities that allow for coarse alignment in specific participant pairs.

**Figure 4.**
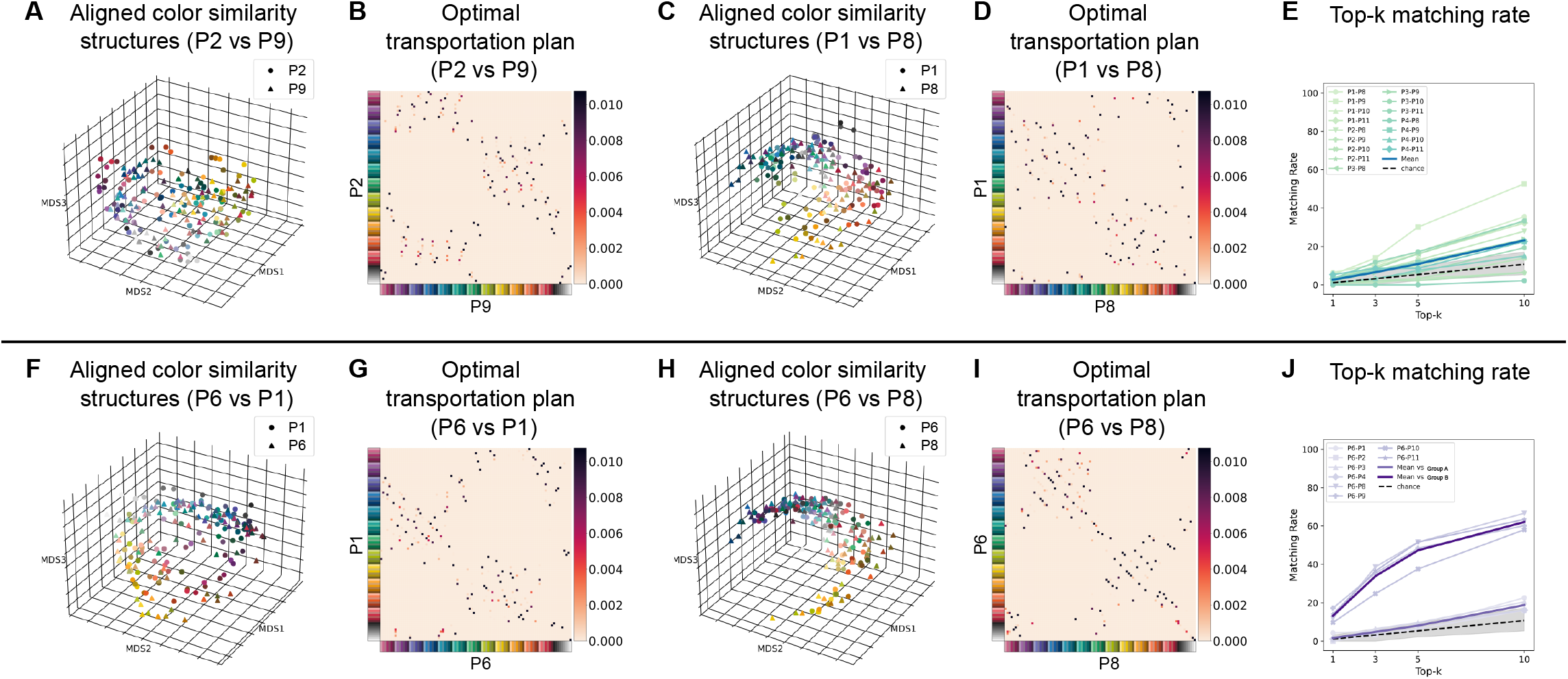
Certain individual pairs fail to align strictly under unsupervised alignment. (A and B, C and D, F and G, and H and I) Aligned color similarity structures of P2 and P9, P1 and P8, P6 and P1, and P6 and P8, respectively, in the same format as Figures 3C and 3D. (E) Top-*k* matching rate for all pairs between Group A (P1–P4) and Group B (P8–P11) (n = 16 pairs). (J) Top-*k* matching rate for all pairs including P6 (n = 10 pairs); mean matching rates for pairs with Group A members and with Group B members are plotted separately. In (E) and (J), the dashed line and gray-shaded region are as in Figure 3F.

### Co-existence of robust groups and individuals who align with neither

Beyond these two groups, we identified a subset of participants (P5, P6, and P7) whose color similarity structures did not robustly align with either group. For example, P6 did not align with a member of Group A (P1: Figures 4F and 4G) or a member of Group B (P8: Figures 4H and 4I), with the optimal transportation plans of each pair showing dispersed probability mass and low matching rates. This pattern is generalized across other pairwise comparisons involving P6 (Figure 4J). Although P6 showed relatively stronger alignment with Group B, these matching rates remained below those observed within either group, thereby indicating that P6 possesses a unique color similarity structure distinct from both groups. Similarly, P5 and P7 did not show robust alignment with either group.

We note that the weaker alignment observed among P5–P7 is unlikely to be attributable to lower measurement precision. To confirm this, we assessed the structural reproducibility of each individual by constructing two color similarity structures from split halves of that participant’s similarity ratings and performing unsupervised alignment between them (see STAR Methods). For every participant, the within-participant matching rate was well above chance, and was comparable to the between-participant matching rates obtained from the full data within each of the two groups we identified before (Figure S4). This indicates that the measurement precision was sufficient for every individual, and that the weaker between-participant alignment of P5–P7 cannot readily be explained by lower within-participant reliability alone.

Combining the results from the previous sections, the full 11×11 individual alignment shows both robustly aligned groups and participants who align with neither of them. To systematically illustrate this global landscape, we examined the structural relationships between individuals through two analytical lenses: the direct quantification of pairwise matching rates (Figure 5A) and a meta-MDS embedding that maps the global structural distances between participants (Figure 5B). In this meta-MDS, each individual is placed in a two-dimensional space such that the distances between points reflect how distant these individuals are, quantified as the difference in the entire histogram patterns separately for *k* =1, 3, 5, and 10 (see STAR Methods). As expected from the preceding analyses, Group A (P1–P4) and Group B (P8– P11) show high peaks within their respective groups (yellow-red and pink-purple bars, respectively, in Figure 5A). In contrast, P5–P7 exhibit intermediate profiles. Specifically, P5 aligns moderately with both groups with slightly better alignment with Group A; P6 aligns more strongly with Group B while remaining distinguishable from it due to lower matching rates; and P7 robustly aligns with neither group, except for its alignment with P8. Consistent with these profiles, P5–P7 are placed between the two groups rather than forming a group of their own (Figure 5B) in the meta-MDS. We refer to them as “structural intermediates” as a descriptive label for this sample. This overall arrangement, i.e., two aligning groups along with three intermediate individuals, was reproduced stably when the meta-MDS was recomputed from random halves of the ratings (Figure S5). The matching rates were lower overall than those obtained from the full data, and accordingly the arrangement was less distinct for the top-1 matching rate. For the top-3, top-5, and top-10 matching rates, however, the same pattern was clearly seen in all 10 partitions.

**Figure 5.**
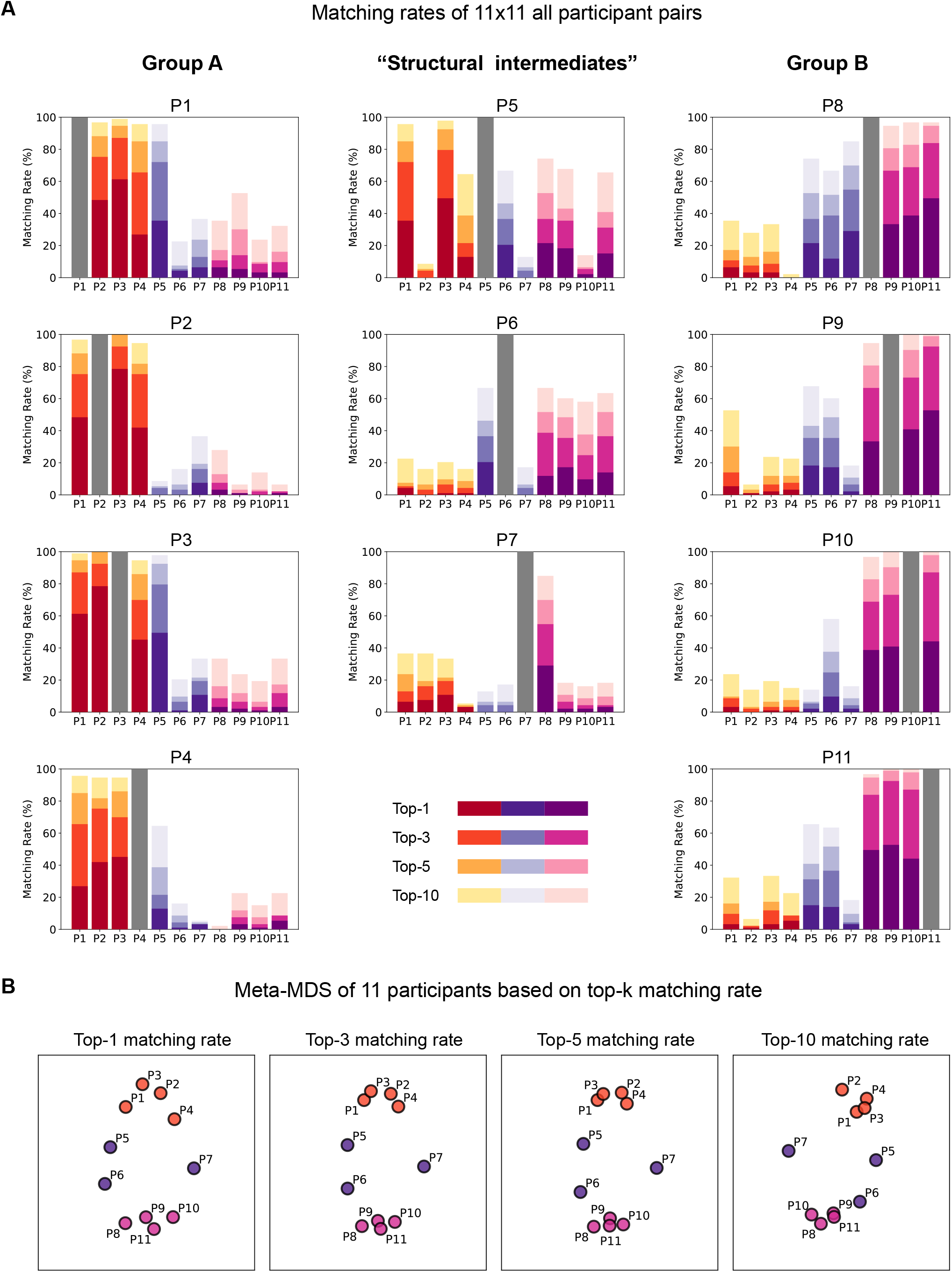
Unsupervised alignment across all 11 participants reveals both robust groups and individuals who align with neither. (A) Matching rates across all 11 × 11 participant pairs. Each panel shows the matching rate of one individual with every other participant. Notably, participants P5–P7, whose FM100 Total Error Scores fell within the range commonly observed among color-neurotypical observers but identified themselves as having atypical color vision, do not align robustly with either Group A (P1–P4) or Group B (P8–P11), and occupy intermediate positions between them in terms of their matching-rate profiles. (B) Meta-MDS of 11 participants based on top-*k* matching rate. Each plot is made by performing 2D MDS using a distance matrix calculated from the top-k matching rate separately for *k* =1, 3, 5, and 10, where distance between participants is computed as the mismatch between histograms (see STAR Methods).

### Correspondence between the structure-based groupings and self-report or FM100 performance

We next examined how the groupings identified above by structural alignment relate to self-report and FM100 performance (Figure 2D; see also Figure S1 and Text S1). All four members of Group A (P1–P4) showed relatively low FM100 Total Error Scores (TES 28), and P1–P3 also self-reported typical color vision. P4, however, self-reported atypical color vision and showed a mild atypical-like error pattern on the FM100. All four members of Group B (P8–P11) showed relatively high Total Error Scores (TES > 100) as well as an atypical (deutan)-like error pattern, and all self-reported atypical color vision. The “structural intermediates” (P5–P7), who aligned with neither group, showed Total Error Scores within the range commonly observed among color-neurotypical observers^39^, yet nonetheless self-reported atypical color vision. Also, P6 and P7 exhibited a moderate atypical (protan)-like error pattern, whereas P5 did not. Note that these correspondences are descriptive rather than diagnostic: determining each participant’s color vision is not the aim of the present study, and the alignment results characterize the structure of color experience rather than color vision itself. None of the three sources of information, namely the alignment results, the FM100, and self-report, is definitive on its own, nor do they always agree.

Collectively, these comparisons highlight what our structure-based approach adds to conventional assessments, and the potential for a structure-based taxonomy of diverse color vision. Our analysis first identified that color similarity structures form two robustly aligned groups, roughly (but not completely) corresponding to color-neurotypicals and atypicals as inferred from self-report and FM100 performance. We note, however, that these should be regarded as candidate groupings in the present sample: additional groups on either side may well emerge with a larger sample. Against this baseline of within-group convergence, the “structural intermediates” (P5–P7) did not fall into either group, and were instead placed between them (Figure 5B). By capturing this fine-grained variation in holistic structure, our unsupervised alignment approach provides a complementary framework for categorizing subjective experiences, revealing individual differences and commonalities from a different angle than the standard color vision metric based on local discrimination sensitivity. Whereas the FM100 asks only whether two colors that look nearly the same can be told apart, our approach looks at how each color relates to every other color: not only to its close neighbors, but also to colors at intermediate distances and to colors at the opposite end of the color space. It then asks whether this global structure of one individual can be mapped onto that of another.

## DISCUSSION

In this study, we demonstrate that color similarity structures can be robustly aligned between individuals in a purely unsupervised manner, and reveal both shared structure and systematic divergence across people. We collected 4,371 pairwise similarity judgments for 93 colors from 11 individuals with diverse forms of color vision. Our primary finding is that structures aligned in a stimulus-matched manner between specific individuals, indicating that my “red” is relationally your “red” and disfavoring the plausibility of structure-preserving swaps such as red-to-green. This was not confined to a single fortunate pair: we identified two groups of individuals, namely Group A and Group B. Within each Group, every pair of similarity structures aligned robustly, while every pair between Groups did not align very well. These groups emerged entirely in a data-driven manner without reference to hue-test performance or self-reported phenomenology, yet post hoc they closely reflected both. Specifically, Group A and Group B corresponded roughly, though not exactly, to color-neurotypical and atypical observers on self-report and the FM100, respectively. At the same time, we also identified certain individuals who did not align with either group (“structural intermediates”). Notably, all of these individuals self-reported atypical color vision despite showing normal color discrimination ability in terms of TES on FM100 test. Together, these results point toward a structure-based taxonomy of color experience grounded in the global structure of reported color similarity, offering a novel categorization that complements standard color vision metrics focused on local sensitivities.

The robust alignment within Group A, whose members were largely color-neurotypical on self-report and the FM100, is not an obvious observation, given the substantial individual differences documented among color-neurotypicals at both neurophysiological and perceptual levels. At the neurophysiological level, color-neurotypicals exhibit considerable variability in their photoreceptors, including a shift in cone spectral sensitivities^40^ and the L:M cone ratio^41,42^. The presence of robust alignment despite these sources of variability suggests that subsequent cortical mechanisms can accommodate such physiological differences^29,43^ and instantiate qualitatively similar color experiences based on a shared structure^44–48^. In fact, P4 exhibited an FM100 error pattern suggestive of mild anomalous trichromacy, a condition in which all three cone types are present but exhibit altered spectral sensitivities (see Figure S1 and Text S1). This observation highlights a potential dissociation between physiological traits and the phenomenal structures captured by our alignment. At the perceptual level, color-neurotypicals also show large and systematic individual differences in the loci of unique hues^49–51^ and in color naming, in which observers cluster into distinct lexical patterns even within the same language^52^. The seeming incompatibility of our results with these findings further suggests that these perceptual traits, including judgments of uniqueness or lexical categorization, may be dissociable from the color qualia themselves.

The robust alignment within Group B, whose members were all atypical on self-report and the FM100, may reflect a contingent outcome rather than a general property of atypical color vision, and should therefore be interpreted carefully. Generally, “atypical” color vision encompasses a diverse range of physiological mechanisms, such as protan or deutan, and varying degrees of severity^35,53–55^. Given this well-documented heterogeneity, the strong within-group convergence seen in Group B in the present sample suggests that these individuals coincidentally share a common physiological subtype. Although we cannot directly determine the precise physiological subtypes of atypical color vision in our dataset, the characteristic error patterns observed in the FM 100 hue test allow for a cautious inference (see Figure S1 and Text S1). Specifically, these patterns are most consistent with deutan-type atypical color vision^39^, in which M-cone function is absent or altered. This inference is also consistent with epidemiological evidence showing that deuteranopia and related deutan conditions are more prevalent than protanopia or tritanopia (involving L-cone or S-cone absence or anomalies, respectively)^53,55^. It should also be noted that the psychophysical data available in the present study do not permit a clear distinction between dichromacy (the complete absence of one cone class) and anomalous trichromacy (the presence of all three cone classes with altered spectral sensitivity)^56^. Such a distinction can in principle be made with an anomaloscope or other more sensitive color discrimination tests, which were not available in this study. It is therefore plausible that, by chance, the individuals in Group B (P8–P11) share relatively similar deutan subtypes or degrees of severity, leading to the observed within-group convergence.

Despite overall structural divergence between the two groups, the presence of coarse alignment in some cross-group pairs (Figures 4C–4E) suggests that aspects of color experiences may still be shared to some extent across these groups. The dominant view based on color models characterizes color-atypical observers as experiencing a restricted subset of neurotypical color qualia. However, a parallel line of work has long suggested that their color experiences can nonetheless be rich and structured in ways comparable to those of neurotypicals^27,57^, consistent with our observations. This view is further supported by a historical fact: in the long history of humanity, “color blindness” was not scientifically recognized until John Dalton’s seminal report in the late 18th century^58^. This delay implies that atypical color vision may have long gone unnoticed because atypical and neurotypical observers communicated without difficulty using a shared color vocabulary grounded in largely overlapping phenomenology. An alternative, less likely but still intriguing possibility is that the shared linguistic system itself shapes color experience in a way that renders their qualia similar^59^.

The “structural intermediates” (P5–P7), which occupy intermediate positions without aligning with either group, may differ from the two groups in the type or degree of color vision atypicality^53–55^. For P6 and P7, their color similarity structures were more similar to those of Group B than to Group A, while still failing to robustly align with either group. This observation is consistent with their FM100 error patterns, which suggest protan-type atypical color vision (see Figure S1 and Text S1), in contrast to the deutan-type patterns that were most consistent with Group B. In addition, their Total Error Scores tended to be lower than those observed in Group B, which aligns with their intermediate position between the two groups. Taken together, these observations suggest that P6 and P7 exhibit moderate, protan-like atypical color vision that is distinct from the pattern observed in Group B. One intriguing possibility is that these participants correspond to anomalous trichromacy, which has been shown to span a continuous spectrum of color vision phenotypes^29,60^, although the FM100 does not allow a reliable distinction between anomalous trichromacy and dichromacy. By contrast, P5 exhibited FM100 performance within the range commonly observed among color-neurotypical observers, despite failing to robustly align with Group A. This pattern leaves open the possibility that P5 exhibits a mild form of color vision variation, such as anomalous trichromacy, that is not detectable by the FM100 but could potentially be revealed by other, more sensitive assessments. How the structural intermediates, as well as the two aligned groups, relate to precisely determined color vision types remains an important subject for future studies.

Our findings highlight the value of a structure-based taxonomy grounded directly in the phenomenology of color experience, beyond what physiological or performance-based diagnoses can provide. Many traditional assessments of color vision, such as the FM 100 hue test and the Ishihara test, often rely on threshold-based performance measures and thus divide observers into coarse categories that may obscure the continuous nature of color vision differences. Moreover, these metrics assess the local discriminability of colors rather than the global structure of the entire color space. Physiological classifications likewise cannot specify how such variations manifest in subjective experience. In contrast, our structure-based classification draws solely on the global color similarity structures inferred from similarity judgments, enabling a phenomenology-first taxonomy of color vision. Notably, our results highlight a dissociation between performance on the FM100 and structure-based classification. For example, P5 exhibited FM100 performance within the normal color discrimination range, yet showed a color similarity structure that was distinct from Group A. Conversely, P4 exhibited an atypical-like error pattern on the FM100 but robustly aligned with Group A. These dissociations indicate that our structure-based approach captures aspects of color experience that are not fully captured by standard performance-based metrics that focus on local sensitivities. One intriguing possibility is to develop structure-based measures that could complement conventional assessments in characterizing individual color vision, although establishing this would require more detailed assessment of each participant’s color vision, such as anomaloscopy, which was not available in this study.

Our one-versus-one alignment framework is inherently generalizable, providing a principled way to capture and compare the structured organization of various aspects of subjective experience across individuals. We and others have already applied this method to multiple domains, including object recognition^61,62^, emotion^63^, and children’s color qualia^31^. The present individual-level validation strengthens the foundation for all these applications. More broadly, the approach is particularly well suited to domains in which individual differences in phenomenal experience are central. These include developmental trajectories (how does qualia structure change from infancy to adulthood?^31^), cross-cultural and cross-linguistic comparisons (do linguistic categories shape qualia structure?^64,65^), and forms of neurodiversity (do individuals with autism or synesthesia organize their experiences differently?^66^). Crucially, the discovery of “structural intermediates” in the present study (individuals whose color similarity structures show idiosyncrasy despite Total Error Scores within the range commonly observed among color-neurotypical observers) suggests that analogous hidden diversity may exist in other perceptual and cognitive domains, with direct implications for clinical assessment and our scientific understanding of human diversity. Ultimately, our approach enables subjectivity itself to be treated within a scientific framework, offering a foundation for addressing how conscious experiences can be shared and compared across minds.

### Limitations of the study

A limitation of our study is the relatively small number of participants (N = 11), which constrains our ability to characterize the full extent of variation in the broader population and to clarify whether this variation is continuous and whether additional groups would emerge. This limited sample size reflects our decision to prioritize the collection of rich datasets employing a substantially larger set of color samples than in previous studies, which used fewer than 30 colors to collect color similarity ratings from each individual^27,29–31,67–70^. This larger set of color samples is necessary to densely sample color space across both hue and tone dimensions, which would not be possible with such small color sets. However, it remains unknown whether additional groups beyond the two identified here would emerge with a larger cohort (as suggested by Palmer^9^). Given the substantial practical burden of obtaining complete sets of similarity judgments for a large number of stimuli, scaling up participant numbers will likely require more efficient sampling methods that can approximate fine-grained similarity structures across a large stimulus space with fewer trials.

## RESOURCE AVAILABILITY

### Lead contact

Requests for further information and resources should be directed to and will be fulfilled by the lead contact, Masafumi Oizumi.

### Materials availability

This study did not generate new materials.

### Data and code availability

- The color similarity judgment data have been deposited at Mendeley Data and are publicly available as of the date of publication at https://doi.org/10.17632/bhx8t6z4ff.1.
- The code for the behavioral experiments has been deposited at Mendeley Data and is publicly available as of the date of publication at https://doi.org/10.17632/bhx8t6z4ff.1. The code used for the analyses in this paper is publicly available at GitHub: https://github.com/oizumi-lab/Individual_Color_GWOT.
- Any additional information required to reanalyze the data reported in this paper is available from the lead contact upon request.

## ACKNOWLEDGMENTS

We thank Yusuke Moriguchi for his valuable comments on the manuscript. M. O. was supported by JST Moonshot R & D Grant Number JPMJMS2012 and Japan Promotion Science, Grant-in-Aid for Transformative Research Areas Grant Number 23H04834. C. H. was supported by Japan Promotion Science, Grant-in-Aid for Transformative Research Areas Grant Number 24H01553. N. T. was supported by National Health Medical Research Council (GNT2037172), Australian Research Council (DP240102680), Japan Society for the Promotion of Science Grant-in-Aid for Transformative Research Areas (A) (23H04830) and Japan Science and Technology (JST) Moonshot R&D Grant (JPMJMS2295), Theoretical Sciences Visiting Program, Okinawa Institute of Science and Technology.

## AUTHOR CONTRIBUTIONS

Y.T., Y.Y., N.T., and M.O. conceptualized and designed the study. Y.T. and Y.Y. conducted behavioral experiments. Y.T. and M.O. performed data analysis. C.H. contributed to the classification and interpretation of color vision test results. Y.T. and M.O. drafted the initial manuscript. All authors reviewed, edited, and approved the final manuscript.

## DECLARATION OF GENERATIVE AI AND AI-ASSISTED TECHNOLOGIES IN THE WRITING PROCESS

During the preparation of this work, the author(s) used ChatGPT, Gemini, and Claude in order to improve the readability and language of the manuscript. After using these tools/services, the author(s) reviewed and edited the content as needed and take(s) full responsibility for the content of the published article.

## SUPPLEMENTAL INFORMATION

Text S1 and Figures S1–S5 with their legends, in a PDF

Video S1. Unsupervised alignment of the four participants in Group A (P1–P4), related to Figure 3.

## STAR⍰METHODS

### KEY RESOURCES TABLE

| REAGENT OR RESOURCE | SOURCE | IDENTIFIER |
| --- | --- | --- |
| Deposited data |  |  |
| Color similarity judgment data | This paper | Mendeley Data: https://doi.org/10.17632/bhx8t6z4ff.1 |
| Software and algorithms |  |  |
| Behavioral experiment code | This paper | Mendeley Data: https://doi.org/10.17632/bhx8t6z4ff.1 |
| Data analysis code | This paper | GitHub: https://github.com/oizumi-lab/Individual_Color_GWOT |

### EXPERIMENTAL MODEL AND STUDY PARTICIPANT DETAILS

#### Participants

Eleven participants (10 males and 1 female; mean ± SD age = 20.9 ± 2.6 years) took part in the experiment. Participants were undergraduate and graduate students from the University of Tokyo. They were recruited via email, which explicitly stated that we were seeking individuals with both typical and atypical color vision. During recruitment, participants self-reported their color vision status: according to these reports, 3 participants had typical color vision (including the female) and 8 had atypical color vision. The experiment was approved by the Research Ethics Committee of the University of Tokyo (approval number: 979-2). All participants provided informed consent prior to the experiment and received monetary compensation upon completion.

## METHOD DETAILS

### Color similarity rating task

#### Apparatus

The experimental stimuli were presented on a VIEWPixx monitor (Vpixx Technologies). The monitor had a width of 52 cm with a resolution of 1920 (W) × 1080 (H) pixels, and the viewing distance was set to 57 cm. The stimuli were created using PsychoPy-2023.2.3^71^. The monitor was calibrated using a ColorCAL MKII Colorimeter (Cambridge Research Systems, Rochester, UK) and the luminance was gamma-corrected. All experiments were conducted in a dark room in which only the monitor provided illumination.

#### Stimuli

93 colors were selected as stimuli from the Practical Color Coordinate System (PCCS) based on those used by Kawakita et al.^24^ (Figure 2B). These 93 colors included 84 chromatic colors, derived from combinations of 12 hues and 7 tones, and 9 achromatic colors. All color stimuli were presented as circles with a diameter of 1 deg. The stimuli were displayed on a background of 1920 × 1080 pixels, each having a randomly assigned RGB value. The mean RGB value of the background pixels was set to [128, 128, 128].

#### Procedure

Each participant rated the similarity of a total of 4,800 color pairs. These included all 4,371 (= _93_ *C*_2_ + 93) possible combinations of 93 colors (including pairs of identical colors), along with an additional 429 randomly selected pairs from the 4,371 pairs (double-pass) to assess response consistency. Each session consisted of similarity ratings for 320 pairs, including 28 or 29 double-pass pairs. The experiment consisted of 15 sessions conducted over five days. The assignment of all color pairs to sessions was randomized for each participant.

In each trial, two color stimuli were presented sequentially on the screen. Participants were then asked to rate the similarity of the two colors (Figure 2A). A fixation point was displayed at the center of the screen for 200 ms, followed by the sequential presentation of two color stimuli for 250 ms each, with an inter-stimulus interval (ISI) of 100 ms. The stimuli were randomly presented at one of four locations, offset by ±1 deg both horizontally and vertically from the fixation point, with the positions of the first and second stimuli adjusted to ensure they did not overlap. The order of color pair presentation was randomized for each participant. A response screen appeared 100 ms after the presentation of the second stimuli. Participants rated the similarity of the two colors by clicking on one of the following values: [-4, -3, -2, -1, 1, 2, 3, 4]. The response scale was explained to participants as follows: [-4: Least similar, -3: Very dissimilar, -2: Dissimilar, -1: Slightly dissimilar, 1: Slightly similar, 2: Similar, 3: Very similar, 4: Most similar]. The scale deliberately omitted a zero response. In our past pilot experiments, having a middle option such as zero unnecessarily induced a strong bias in some participants, though not all of them, and some participants explicitly reported resorting to the middle response option as they did not want to report something subtle. Participants were instructed to use the full range of the scale, including intermediate values, when making their ratings. After responding, participants were asked to click on a small square in the center of the screen so that they fixate the center of the cross in the next trial.

At the beginning of each session, participants completed 20 practice trials to remind themselves of the task procedure, recalibrate the similarity ratings, and adapt to the luminance level of the experimental room. During the practice, participants received feedback on their responses, which was limited to which rating they used and what the particular rating means. The results of the practice trials were excluded from analysis. During the main trials, a pause screen was displayed every 40 trials, allowing participants to take short breaks as needed. Each session took approximately 20 minutes to complete.

To evaluate the internal consistency of participants’ responses, we computed a double-pass correlation for each participant by pooling all stimulus pairs that were presented twice across sessions. The double-pass correlation was defined as the Pearson correlation coefficient between the two sets of ratings given to the same stimulus pairs. This measure reflects the degree to which a participant consistently judged the similarity of identical color pairs across repeated presentations. Across participants, the number of double-pass pairs was *N* = 429, yielding a degree of freedom DOF = 427 for each correlation estimate. The results showed high consistency across all participants, with correlation coefficients ranging from *r* = 0.65 to *r* = 0.88 (mean *r* = 0.79). All correlations were highly significant (all *p* < 10^-50^), indicating that the similarity judgments were reliable and stable within individuals. Session-by-session double-pass correlations were broadly consistent with the corresponding across-session correlations, although some within-participant variability was observed (Figure S3). One session from participant P1 showed a near-zero correlation and appeared as a clear outlier relative to other sessions across participants. Still, we decided not to exclude this session, as no session-level exclusion criterion based on double-pass correlation had been specified a priori.

### Color vision assessment by FM 100 hue test

#### Apparatus

The Farnsworth-Munsell 100 Hue Test (X-Rite) was employed. All experiments were conducted in a dark room, and the Pantone 3 light booth (X-Rite) was used as the light source device.

#### Stimuli

The test utilized 85 physical colored caps, which were divided into four trays. Each tray contained 22 or 23 caps arranged in random order.

#### Procedure

Participants were instructed to arrange the caps in each tray in the order of their hues. The sorting task was conducted under a standard daylight light source (D65) provided by the light booth. Although the original FM 100 Hue Test specifies Illuminant C as the standard light source, the difference between Illuminant C and D65 has been shown to have a negligible effect on FM 100 Hue Test performance^72^. Participants performed two sessions in total, each consisting of one sorting task per tray (four trays in total). The time required to sort one tray was estimated at approximately two minutes, but no time limit was imposed for the task.

#### Analysis

Each colored cap was numbered sequentially from 1 to 85 based on its hue. For each cap, a score was calculated as the sum of the absolute differences in hue number between the cap and its adjacent caps. For example, if the caps were arranged in the order 10, 11, 12, 16, and 14, the score for cap 11 would be ∥ 10 – 11 ∥ + ∥ 12 – 11 ∥ ∥ = 2, and the score for cap 16 would be ∥ 12 – 16 ∥ + ∥ 14 - 16 ∥ 1= 6. For each cap, the adjusted score was calculated by subtracting 2 from its original score, and the total sum of these adjusted scores across all caps was calculated as the Total Error Score (TES). In addition to this summary measure, cap-wise error scores were examined to characterize the distribution of errors across hues (Figure S1). Based on the error patterns, the subtype of atypical color vision was inferred according to the error patterns depicted by Farnsworth^39^ (see Text S1).

### QUANTIFICATION AND STATISTICAL ANALYSIS

#### Construction of color similarity structures

First, we estimated the color similarity structure of each participant using the similarity ratings obtained from the psychophysics experiment (Figure 2A). Assuming that the distance relationships between color experiences can be modeled within a Euclidean space, we estimated the color embeddings for each participant using Multidimensional Scaling (MDS) implemented in scikit-learn^73^. Specifically, for the *i*-th color (*i* ε 1,2,…,93), the embedding *x*_*i*_ in a 5-dimensional Euclidean space was computed by minimizing the Stress value defined by the following equation:

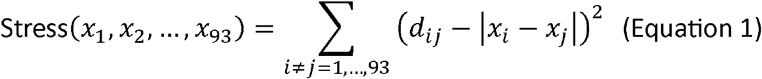

Here, *d*_*ij*_ denotes the dissimilarity rating between stimuli *i* and *j*, while |*x*_*i*_*-x*_*j*_| denotes the Euclidean distance between the embeddings *x*_*i*_ and *x*_*j*_. The dissimilarity values *d*_*ij*_ were obtained by converting the raw similarity ratings in the range of [-4,-3,-2,-1,+1,+2,+3,+4] into integer scores in the range of [7,6,5,4,3,2,1,0] in this order, where higher values indicate greater dissimilarity. For example, the rating of -4 (“least similar”) was converted to 7, and the rating of +1 (“slightly similar”) was converted to 3. This conversion, which assigns equally spaced values to the ordered response categories, is justified on the following grounds. In our past pilot experiments, almost all participants reported a feeling of unit intervals with this scale. And we observed no difference in the resulting pattern when we used a 0–7 rating scheme and a -4 to -1 and +1 to +4 scheme, except that 0–7 rating confused some participants leading to reverse response (e.g., 0 being ‘most different’)^30^. For one participant (P8), a single response value (one of all possible combinations of colors) was missing, and this missing dissimilarity rating was imputed with a neutral value of 3. For color pairs included in the double-pass trials, we averaged the ratings from the first and the second presentations. This process corresponds to finding the embeddings of all colors such that the distances between them approximate the dissimilarity ratings as closely as possible.

In all subsequent analyses, we used the estimated distance structures rather than the raw similarity ratings. For each participant, 93 × 93 color dissimilarity matrix *D* was computed based on the Euclidean distances between the embedded colors, where each element *D*_*ij*_ represents the subjective dissimilarity between the *i*-th and *j*-th color experiences: *D*_*ij*_= |*x*_*i*_*-x*_*j*_|. Although we obtained similarity ratings for all possible color pairs, we relied on the distances derived from the estimated color similarity structures for two reasons. First, we assume that the dissimilarities between color experiences are inherently continuous within each participant, whereas the collected ratings were discretized by the response format. Second, since each color pair was rated only once (except for the double-pass trials), occasional response errors (e.g., accidental button presses) could occur. However, such local noise would be effectively attenuated through the embedding procedure, which aggregates information across all pairwise ratings to find a globally consistent geometric configuration.

For visualization, we applied MDS (scikit-learn^73^) to the distance structure of the 5-dimensional embeddings and retained the first three dimensions to obtain a lower-dimensional representation of the color similarity structure.

#### Gromov-Wasserstein Optimal Transport (GWOT)

To evaluate the structural correspondence between two individuals’ color similarity structures, we employed Gromov–Wasserstein Optimal Transport (GWOT). GWOT provides an unsupervised framework for identifying the maximally structure-preserving mapping between two metric spaces, in this case, two participants’ color similarity structures, denoted by color dissimilarity matrices *D* and *D*′. The correspondence is obtained by minimizing the Gromov-Wasserstein Distance (GWD), which quantifies the discrepancy between pairwise distance relationships across the two spaces:

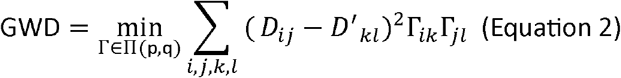

Here, Π(p,q) denotes the feasible set of transportation plans Γ, such that Γ_*ik*_ ≥ 0, ∑_*k*_Γ_*ik*_ = *p*_*i*_, and ∑_*i*_ Γ_*ik*_ = *q*_*k*_, where p and q are the marginal distributions over the two spaces. In our setting, both marginals were uniform, i.e., *p*_*i*_ = *q*_*k*_ = 1/93 for all *i,k*, imposing an equal-mass correspondence between all colors.

Because the optimization problem is non-convex, multiple local minima arise depending on initialization. To efficiently search for a good local optimum, we add an entropy-regularization term to the objective function of GWOT as follows:

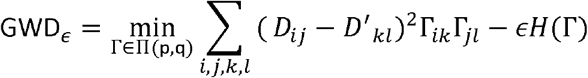

where *H*(Γ) is the entropy of a transportation plan Γ and ε is a hyperparameter that determines the strength of the entropy regularization.

To approximate the global optimum, we performed the optimization 500 times with random initializations of Γ and different ε values for each participant pair and picked the solution yielding the lowest GWD value without the entropy-regularization term (Equation 2). Each element in the initial transportation plan was sampled from the uniform distribution [0, 1], and normalized to satisfy the condition of the feasible set of transportation plans Π(p,q). We sampled 500 different values of ε ranging from 0.01 to 0.1 using the Tree-structured Parzen Estimator (TPE) sampler^74^. Prior to applying GWOT, the two compared dissimilarity matrices were histogram-matched to equalize their overall scale, because GWOT evaluates structural correspondence only up to isometry and is, therefore, sensitive to differences in scale between the two spaces. This optimization was implemented using the Gromov-Wasserstein Optimal Transport Toolbox (GWTune)^26^.

#### Evaluation of the unsupervised alignment

To quantify the degree to which two individuals’ color similarity structures were aligned, we computed the Matching Rate between two structures. Matching Rate represents the proportion of colors that were mapped to the same physical color under the optimal transportation plan Γ obtained by GWOT. To prepare for the computation of Matching Rate, we denote the color labels in participant 1 and 2 as *c*_1_ and *c*_2_, respectively. For each color *i* in participant 1, denoted by *c*_1i_, the matching condition is defined as:

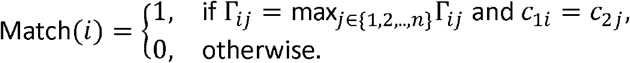

This function indicates whether the *i*-th color in participant 1 corresponds to the same physical color in participant 2 under the optimal mapping. The Matching Rate is then given by the proportion of colors in participant 1 that match with the same physical colors in participant 2:

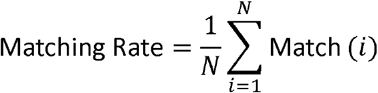

where N(= 93) is the total number of colors. A higher Matching Rate indicates stronger structural alignment between the two color similarity structures, whereas a lower value suggests greater individual divergence.

The matching rate defined above is the top-1 matching rate. More generally, we also define top-*k* matching rate, which evaluates more coarse alignment between two structures. For each color *i* in participant 1, we can define a function to determine if the probability of the *i*-th color corresponding to the same color in participant 2 is within the top-k probabilities:

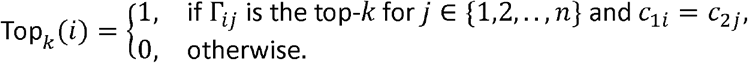

The top-*k* matching rate was calculated as

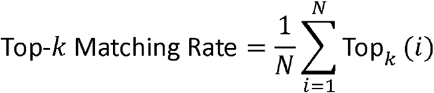

The chance-level matching rate was estimated by randomly generating 10,000 matrices satisfying the constraints of a transportation plan. For each matrix, the matching rate was computed, yielding a null distribution of matching rates. Finally, the 2.5th–97.5th percentiles of this distribution were taken as the 95% percentile interval for the chance-level matching rate.

#### Visualization of the unsupervised alignment

We visualized the results of the unsupervised alignment to intuitively demonstrate how the color similarity structures from different participants are aligned in a shared space. Here, we used the color embeddings *X* and *Y* and the optimal transportation plan Γ that had been obtained in the previous analysis steps. Each embedding matrix *X,Y* ∈ ℝ^*d*×*n*^ (with *d* = 5, *n* = 93) represents the color similarity structure for one participant, where each column corresponds to a color and each row corresponds to one embedding dimension. Thus, the distances among columns in *X* and *Y* reflect the perceptual dissimilarity between colors estimated from individual similarity judgments. Multiplying *Y* by the optimal transportation plan Γ from the right (i.e., *Y*Γ) yields a transformed embedding in which each column of *Y*Γ represents a weighted combination of the original embeddings in *Y*, according to the correspondence encoded in Γ. In the special case where Γ is a permutation matrix, this transformation reduces to a simple reordering of the columns of *Y* in accordance with the one-to-one correspondence specified by Γ.

To align the two structures *X,Y* in the same coordinate system, we searched for an orthogonal transformation matrix *Q* that rotates and reflects *Y*Γ so that its overall geometric configuration best overlaps with *X*. This motivation leads to the following orthogonal Procrustes problem:

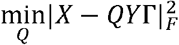

where |·|_*F*_ denotes the Frobenius norm. The optimal *Q* was obtained via the singular value decomposition (SVD) of *X*(*Y*Γ)^T^. The aligned color embeddings were then visualized in a common space using *X* and *QY*Γ.

#### Split-half reliability analysis

To estimate how precisely each participant’s color similarity structure was measured, we conducted a within-participant split-half reliability analysis. The 4,371 color pairs used to construct each individual’s color similarity structure were randomly partitioned into two disjoint, equal-sized halves, independently of session structure. Because each half covers only about half of the pairs, Equation 1 cannot be applied directly. We therefore obtained a 5-dimensional embedding *X* ∈ *ℝ*^93×5^ for each half using a weighted variant of MDS, implemented in Python following the SMACOF algorithm^75^, which accommodates partially observed dissimilarity ratings by assigning a weight to each pair, by minimizing σ(*X*) defined as below:

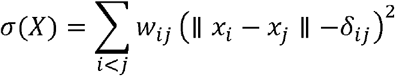

Here, *δ*_*ij*_ is the dissimilarity rating between stimuli *i* and *j, x*_*i*_ is the embedding of the *i*-th color, and *W*_*ij*_ is set to 1 for pairs contained in the half and to 0 otherwise, so that unobserved pairs are excluded from the stress function. With all pairs observed, this reduces to Equation 1.

A 93 x 93 dissimilarity matrix was then computed from the Euclidean distances between the embeddings of each half, and the two matrices were aligned using GWOT with the same settings as in the between-participant analysis. The resulting top-*k* matching rate provides an estimate of how well each participant’s own structure reproduces itself under the measurement noise present in the data. We repeated the whole procedure for 10 independent random partitions per participant. The results are shown in Figure S4.

We also assessed the stability of the groupings using the same split halves. For each partition, we estimated each participant’s color similarity structure from one half of the ratings, applied GWOT to all pairs of participants, and recomputed the meta-MDS embedding from the resulting top-k matching rates. This was repeated for the same 10 partitions, and the results are shown in Figure S5.

#### Meta-MDS of 11 participants based on top-k matching rate

To visualize how the 11 individuals resemble each other or diverge in their color similarity structures, we embedded participants into a two-dimensional space using the following procedure. First, for each participant, we computed a histogram of the top-*k* matching rates obtained from pairwise alignments with all participants, where the self-matching rate was fixed at 100% (as shown in Figure 5A). The dissimilarity between a pair of participants was then defined as the *L*_1_ distance between their corresponding top-*k* matching-rate histograms; that is, we subtracted one histogram from the other and summed the absolute values of the resulting differences. Using the resulting participant-by-participant distance matrix, we performed two-dimensional multidimensional scaling (MDS) and plotted each participant in the resulting embedding space. This entire procedure was performed separately for *k* = 1,3,5, and 10.

