## Supplementary material for "Individual-to-individual mapping of color similarity structures: Is my “red” relationally your “red”?": Text S1 Figures S1-S5

### Supplemental information

#### This PDF file includes:

Text S1
Figures S1–S5

#### Other supplemental information for this manuscript:

Video S1

**Text S1. Classification and interpretation of FM100 hue test results**

We inferred both the presence of atypical color vision and the subtype of atypicality based on cap-wise error patterns described by Farnsworth[39] for the Farnsworth–Munsell 100 Hue Test (FM100). We emphasize at the outset that these inferences are not intended as definitive categorizations of underlying physiological subtypes, but rather as descriptive classifications based on characteristic error patterns observed in the FM100. To infer the color vision subtype, we examined the distribution of cap-wise errors in the circular plot. Although errors were present across the full circle, we focused primarily on the right half of the plot (cap numbers 44–85), where the protan and deutan confusion axes are most clearly differentiated, whereas these axes are substantially closer in the left half and therefore less informative for subtype discrimination. Specifically, when the dominant error peak overlapped with the protan confusion axis but not with the deutan confusion axis, we inferred protan-type atypical color vision. When the dominant error peak overlapped with the deutan confusion axis, we inferred deutan-type atypical color vision, regardless of any concurrent overlap with the protan axis. Participants who did not show a clear overlap with either confusion axis were inferred to be color-neurotypical.

Applying these criteria, we inferred participants P8–P11 to exhibit deutan-type atypical color vision, whereas P6 and P7 were inferred to exhibit protan-type atypical color vision. Although the Total Error Score (TES) of P4 fell within the range commonly observed among color-neurotypical observers, this participant showed a consistent, axis-specific error pattern. We therefore classified P4 as exhibiting a mild form of atypical color vision with a protan- or deutan-like error pattern. Given the mild nature of this pattern, one possible interpretation is anomalous trichromacy; however, this cannot be confirmed, as the FM100 does not allow a reliable distinction between anomalous trichromacy and dichromacy. The remaining participants (P1, P2, P3, and P5) were inferred to be color-neurotypical, as their error scores fell within the normal range and they did not show axis-specific error patterns characteristic of atypical color vision.

Importantly, these subtype assignments should be regarded as cautious inferences based on characteristic error patterns, rather than definitive physiological diagnoses. In particular, the FM100 does not permit a definitive distinction between protan and deutan subtypes in all cases, and our inferences are therefore intended to be descriptive rather than diagnostic.

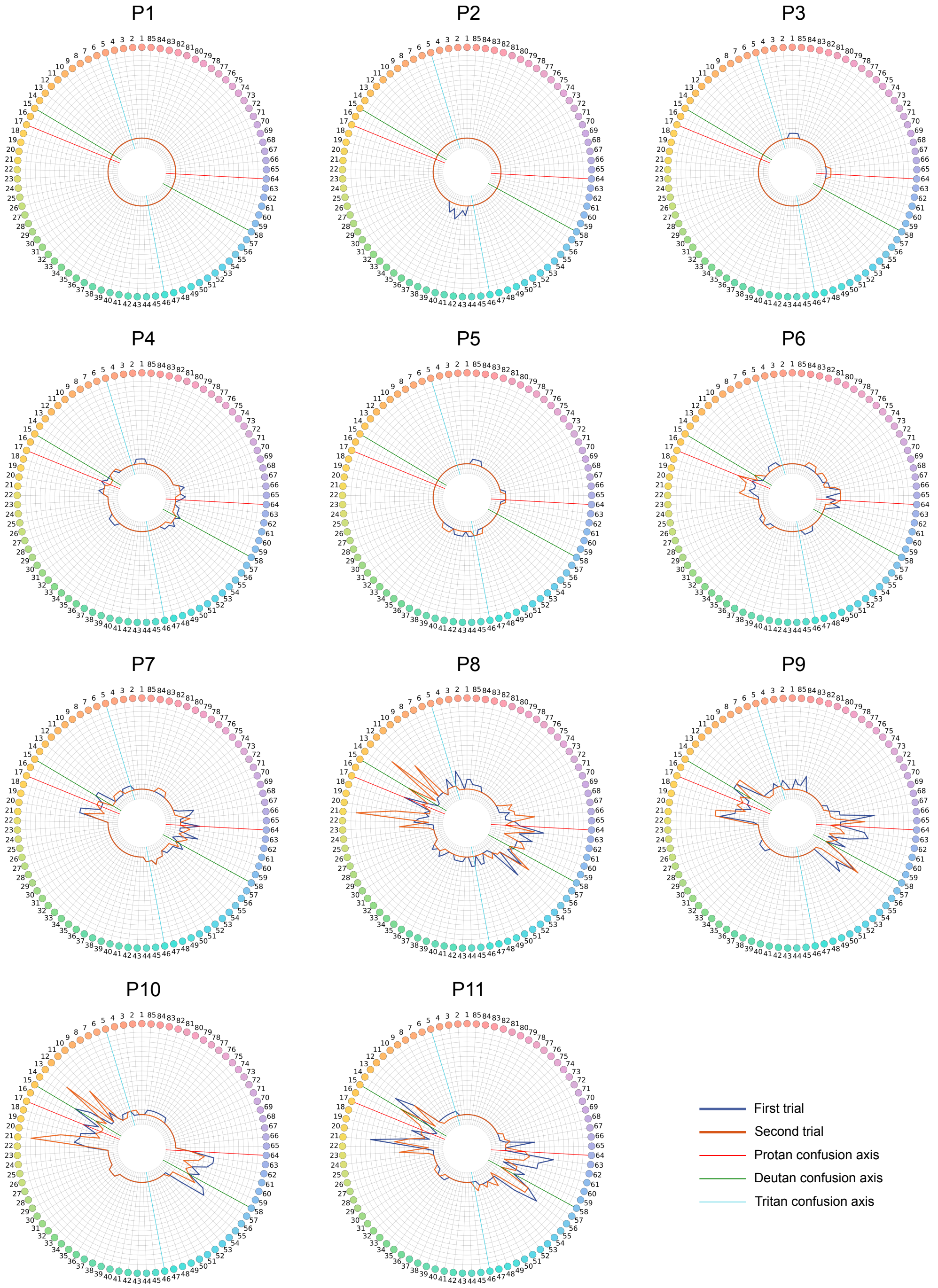


**Figure S1. Error patterns on the FM100 Hue Test for 11 participants**
Each colored circle indicates the color of a cap, and the number attached to each circle denotes the cap number. Blue and orange lines show the per-cap error scores in the first and second trials, respectively (minimum score = 2). Red, green, and sky-blue lines indicate the protan, deutan, and tritan confusion axes, respectively.

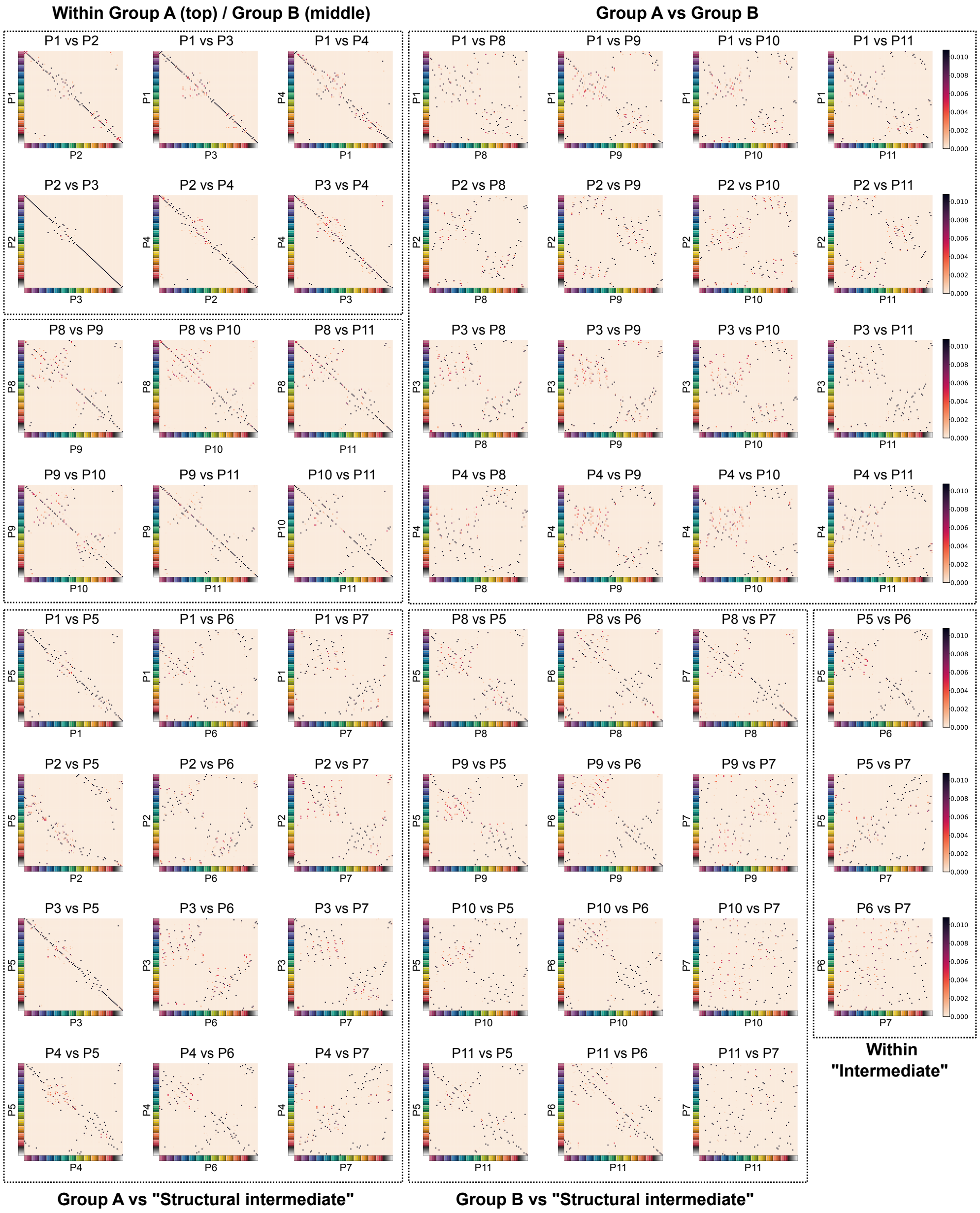


**Figure S2. Unsupervised alignment of all 11 × 11 participant pairs**
Each matrix shows the optimal transportation plan, in the same format as Figure 3D.

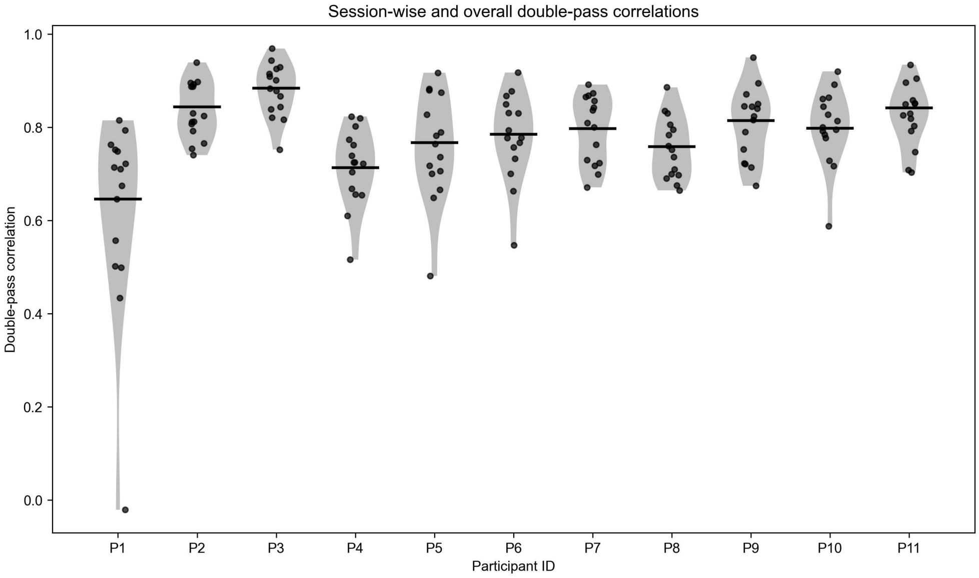


**Figure S3. Double-pass correlations for 11 participants**
Each point represents the correlation computed from double-pass pairs within a single session. Horizontal bars indicate the across-session double-pass correlations computed by pooling all double-pass pairs across sessions.

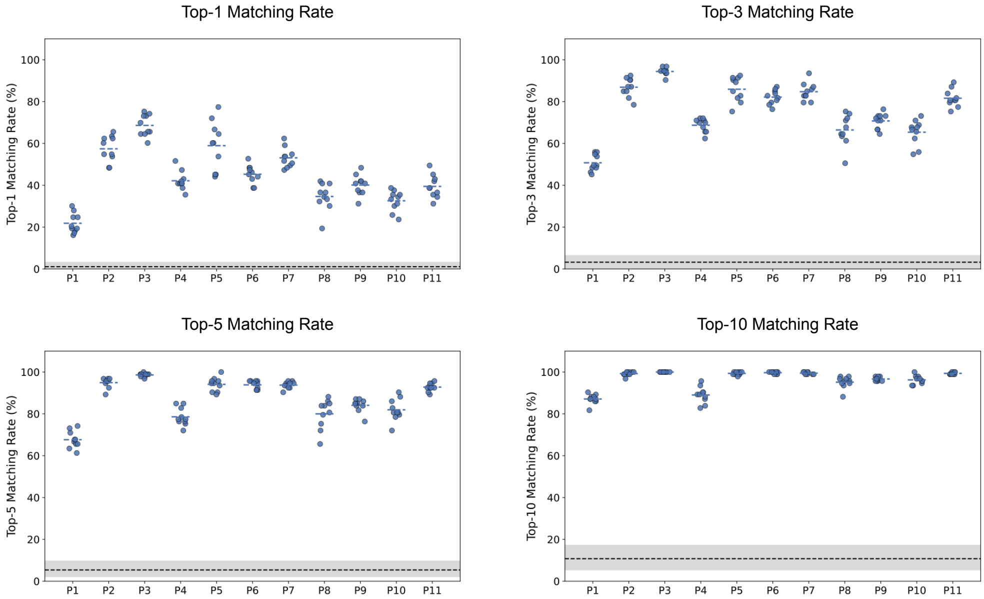


**Figure S4. Within-participant split-half reliability of individual color similarity structures**

Each panel shows the top-$k$ matching rate between the two half-structures for $k = 1, 3, 5, 10$, for each individual. Each point corresponds to one of 10 independent random partitions, and the horizontal bar indicates the mean across partitions. The black dashed line denotes the mean chance-level matching rate, and the shaded band denotes the 95% percentile interval of the chance-level distribution.

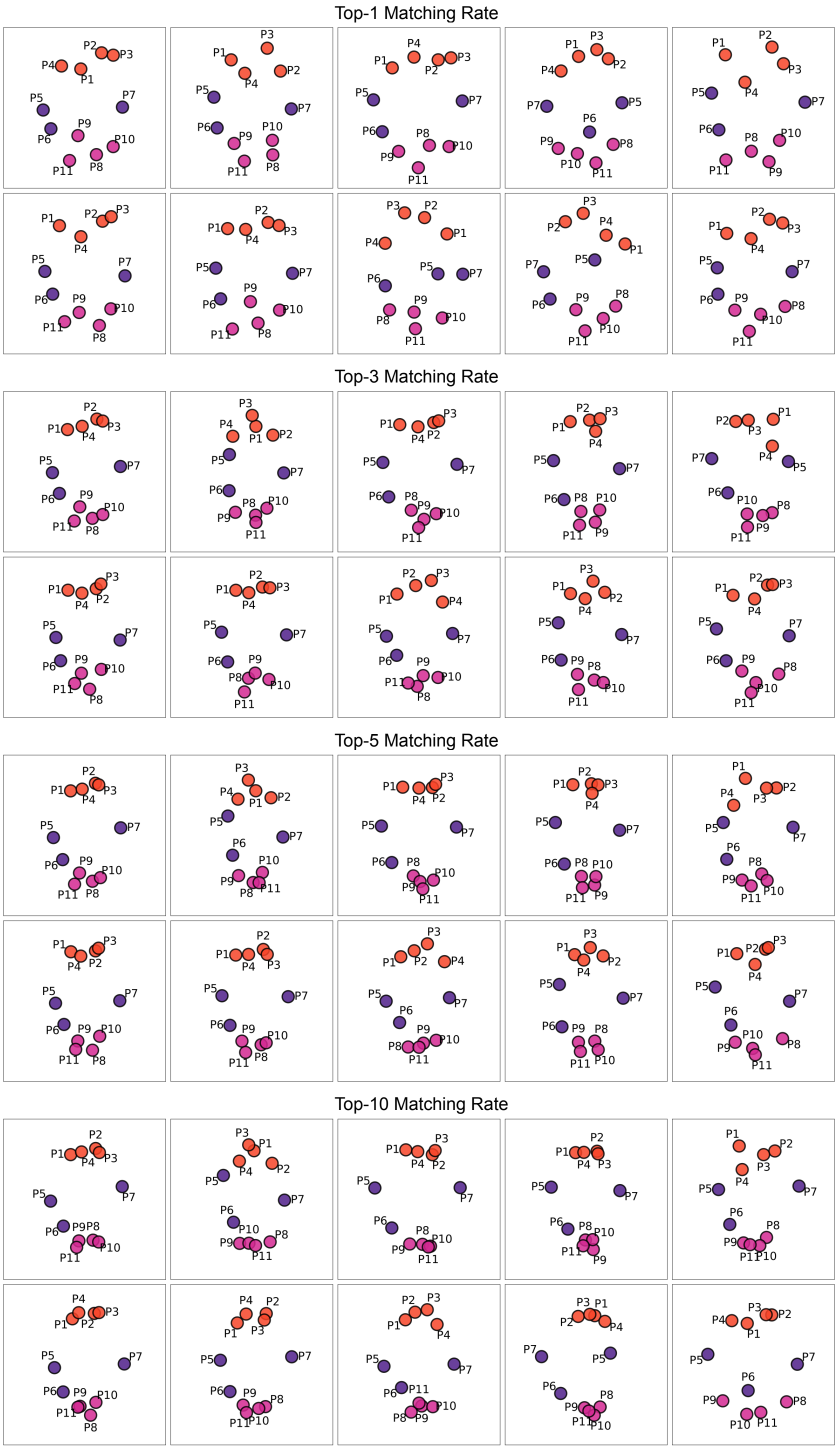


**Figure S5. Stability of the groupings across resampled trials**

Each panel shows the meta-MDS embedding of the 11 participants computed from one random half of the ratings, in the same format as Figure 5B. Ten independent partitions are shown for each of the top-$k$ matching rates ($k = 1, 3, 5, 10$).
